# Closed-loop electrical stimulation to prevent focal epilepsy progression and long-term memory impairment

**DOI:** 10.1101/2024.02.09.579660

**Authors:** Jose J. Ferrero, Ahnaf R. Hassan, Zelin Yu, Zifang Zhao, Liang Ma, Cynthia Wu, Shan Shao, Takeshi Kawano, Judah Engel, Werner Doyle, Orrin Devinsky, Dion Khodagholy, Jennifer N. Gelinas

**Affiliations:** Department of Neurology, Columbia University Irving Medical Center, New York, USA; Department of Biomedical Engineering, Columbia University, New York, USA; Department of Electrical Engineering, Columbia University, New York, USA; Department of Electrical and Electronic Information Engineering, Toyohashi University of Technology, Toyohashi, Japan; Institute for Research on Next-generation Semiconductor and Sensing Science (IRES2), Toyohashi University of Technology, Toyohashi, Japan; Comprehensive Epilepsy Center, New York University, New York, New York, USA

**Author notes:** These authors contributed equally (Jose J. Ferrero, Ahnaf R. Hassan). Corresponding authors: Dion Khodagholy Jennifer Gelinas.

**Keywords:** epilepsy, oscillations, closed-loop stimulation, memory

## Abstract

Interictal epileptiform discharges (IEDs) are ubiquitously expressed in epileptic networks and disrupt cognitive functions. It is unclear whether addressing IED-induced dysfunction could improve epilepsy outcomes as most therapeutics target seizures. We show in a model of progressive hippocampal epilepsy that IEDs produce pathological oscillatory coupling which is associated with prolonged, hypersynchronous neural spiking in synaptically connected cortex and expands the brain territory capable of generating IEDs. A similar relationship between IED-mediated oscillatory coupling and temporal organization of IEDs across brain regions was identified in human subjects with refractory focal epilepsy. Spatiotemporally targeted closed-loop electrical stimulation triggered on hippocampal IED occurrence eliminated the abnormal cortical activity patterns, preventing spread of the epileptic network and ameliorating long-term spatial memory deficits in rodents. These findings suggest that stimulation-based network interventions that normalize interictal dynamics may be an effective treatment of epilepsy and its comorbidities, with a low barrier to clinical translation.

**One-Sentence Summary:** Targeted closed-loop electrical stimulation prevents spread of the epileptic network and ameliorates long-term spatial memory deficits.

## Introduction

Focal epilepsies are associated with large-scale structural and functional neural network abnormalities that can extend beyond the brain regions responsible for seizure generation(*1*). These derangements are associated with neuropsychiatric comorbidities that can worsen over time(*2*). Epilepsy therapeutics focused on eliminating seizures have had limited efficacy in modifying disease course and addressing these comorbidities, which can profoundly impair quality of life(*3*–*5*). Epileptic networks predominantly exist in the interictal state, which contains aberrant dynamics and epileptiform patterns that interfere with physiological processes(*6*, *7*). However, it remains unclear whether and how altering interictal epileptiform activity could treat epilepsy and its comorbidities.

Continuous, intermittent deep brain stimulation (DBS) and vagus nerve stimulation (VNS) are aimed to impact hubs of highly interconnected, complex networks in an empiric attempt to induce desynchronization and prevent seizure initiation(*8*, *9*). Closed-loop electrical stimulation (such as Responsive Neurostimulation Systems; NeuroPace Inc. RNS System) can be configured to deliver abortive stimulation to the seizure onset zone in response to detected ictal patterns(*10–13*). In each case, seizure reductions emerge well after therapy onset, implicating chronic, plasticity-related mechanisms(*14*, *15*). Neuropsychiatric outcomes of these approaches in patients with epilepsy vary(*16*), but efficacy in primary psychiatric patients indicates the potential tractability of such symptoms to interictal network modification(*17*, *18*). Further, in animal epilepsy models, interventions that normalize interictal state dynamics can improve memory(*19*). These results support that mechanistically driven manipulation of interictal patterns could gradually reshape neural networks to support physiological brain functions and suppress epileptic activity.

The potential target mechanisms to achieve these goals in epileptic networks are mostly unknown, but could include amelioration of deranged physiological interactions. During non-rapid eye movement (NREM) sleep, consolidation of episodic memory requires precise correlation of hippocampal and cortical oscillations including hippocampal sharp wave-ripples, the cortical slow oscillation, cortical spindles, and cortical ripples(*20–23*). Interictal epileptiform discharges (IEDs), a key pathological output of the interictal state, disrupt these critical interactions by initiating strong, precise temporal coupling with sleep spindles which surpasses physiological ripple-spindle correlation(*24*). IED-spindle coupling occurs in rodent models and human patients with focal epilepsy, establishing this phenomenon as a potential interictal therapeutic target(*25–27*).

Here we show that this IED-spindle coupling drives prolonged, hypersynchronous cortical spiking and predisposes to generation of local cortical IEDs, effectively spreading the epileptic network. Our data support that a similar process may establish independent foci of interictal epileptiform activity in patients with focal epilepsy. Cortical closed-loop electrical stimulation that inhibits IED-spindle coupling can prevent expression of cortical IEDs, mitigating enlargement of the epileptic network, and preserve long-term memory in focal epilepsy. These results support the use of spatiotemporally focused, interictal-based approaches to normalize epileptic activity patterns and address neuropsychiatric comorbidities.

### Hippocampal IEDs can create an independent cortical interictal focus

To examine how hippocampal-cortical network interactions were altered in the presence of ongoing, progressive epileptic activity, we used hippocampal kindling in freely moving rats while acquiring *in vivo* electrophysiology data from hippocampus and a key synaptically connected cortical area, medial prefrontal cortex (mPFC). We extended an established protocol(*24*) to permit more rapid transition from focal hippocampal to bilateral convulsive seizures, with rats consistently advancing to Racine stage 4 after 10-15 days of kindling (**Figure 1a, Supplementary Figure 1a**). During NREM sleep, we observed a shift in the coherence of hippocampal-mPFC activity. Early kindling (days 5-10) was associated with a broad frequency spectrum increase in coherence, driven prominently by hippocampal IED-induced sharp transients in the mPFC. With further progression of kindling (late kindling, days 15-20), coherence decreased and spontaneous, hippocampal-independent waveforms meeting criteria for IEDs became apparent in the mPFC (**Figure 1b-c**). Hippocampal IED occurrence rate demonstrated an early, rapid increase whereas mPFC IEDs emerged at a delay, with occurrence accelerating only late in kindling (**Figure 1d; Supplementary Figure 1b**). Concordant with coherence analyses, we found that mPFC IEDs were strongly linked to hippocampal IEDs in early kindling, but subsequently a subset became independent (**Figure 1e**). Hippocampal spectrograms trigger-averaged on occurrence time of mPFC IEDs demonstrated a broadband, high power transient in the hippocampal LFP, consistent with a co-occurring hippocampal IED in early kindling (**Supplementary Figure 1c, upper**). However, this power significantly decreased during late kindling, indicating that mPFC IEDs were now occurring independently as well (**Supplementary Figure 1c, lower**). Furthermore, cross correlation of IEDs detected separately in hippocampus and mPFC revealed a highly significant peak of hippocampal IED occurrence immediately preceding the mPFC IED in early kindling that decreased as kindling progressed (**Supplementary Figure 1d**). Independent mPFC IEDs (occurring > 100 ms from a hippocampal IED) were detected in all rats, with incidence higher at the later compared to earlier kindling stages (**Figure 1f**). These results indicate that progression of epileptic activity in the hippocampus reliably leads to expression of IEDs in the mPFC. These mPFC IEDs eventually lose coherence with hippocampal IEDs, suggesting that the mPFC has become an independent focus of interictal epileptic activity.

**Figure 1.**
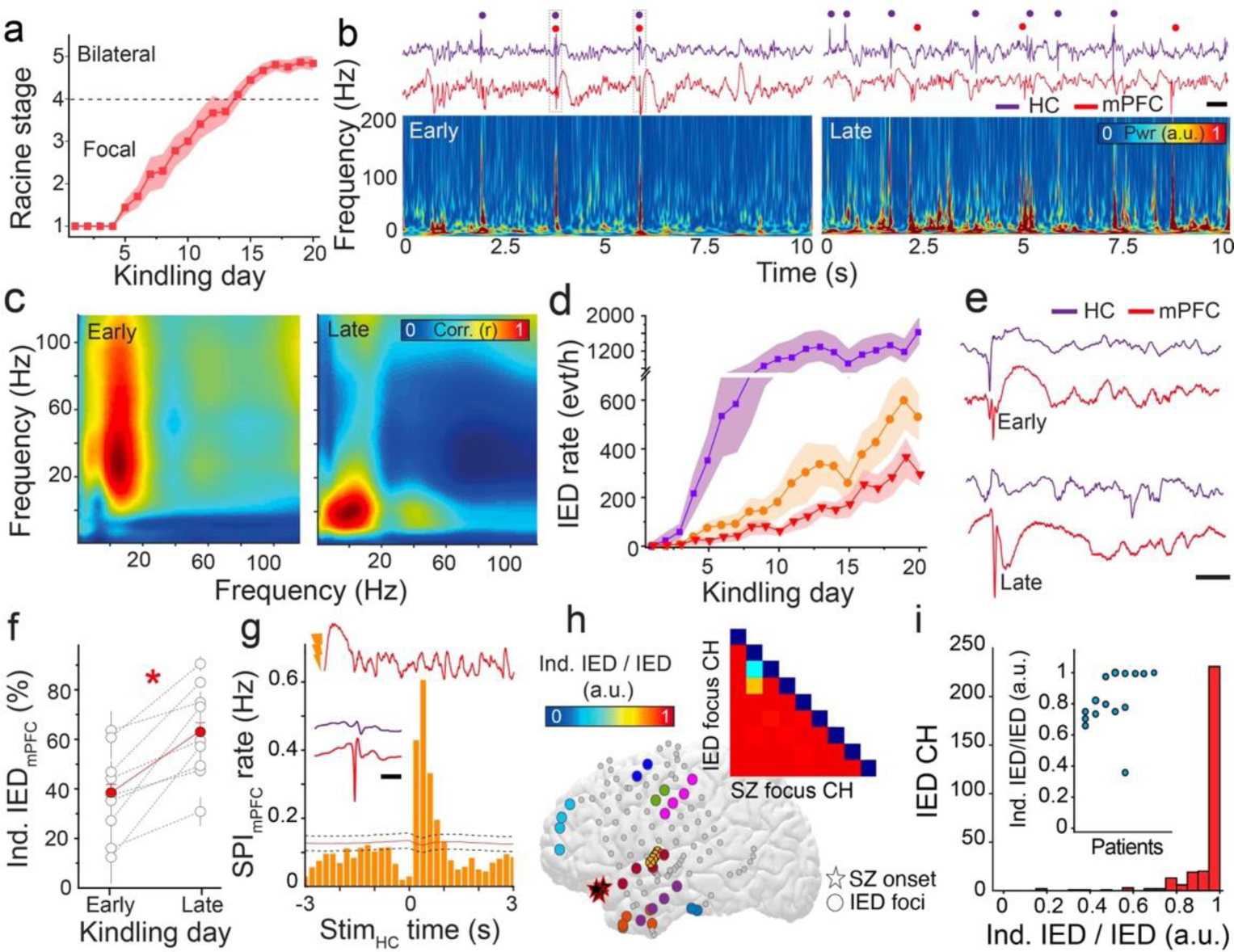
Progression of focal epilepsy is associated with creation of an independent focus of interictal epilepsy activity. **(a)** Average Racine stage progression across kindling (n = 10 rats; shaded error bars represent SEM). **(b)** Sample wide-band LFP traces acquired simultaneously from hippocampus (HC, purple) and mPFC (red; upper), with corresponding mPFC spectrogram (lower) for early stage (days 5 to 10) and late stage (days 15 to 20) of kindling; dots indicate time of IED occurrence. Scale bar: 500 ms. **(c)** Sample comodulograms demonstrating decreased cross-frequency coupling between hippocampus and mPFC during progression from early (left) to late stage (right) of kindling. **(d)** Occurrence of IEDs in hippocampus (purple) and mPFC (total, orange; independent, red) in NREM sleep over kindling (n = 10 rats; shaded error bars represent SEM). **(e)** Sample raw LFP from hippocampus (HC, purple) and mPFC (red), showing representative mPFC IED coupled with hippocampal IED (upper) and independent mPFC IED (lower). Scale bar: 100 ms. **(f)** Increase in the percentage of independent mPFC IEDs from early to late stage of kindling (n = 10 rats, t = - 5.34, P = 1.03 × 10 ^-9^). **(g)** Electrical pulses to the hippocampal commissure generate a-IED_HC_ that induce spindles and independent IEDs in mPFC. Representative raw mPFC trace after pulse stimulation (upper), average independent mPFC IED trace (red) with corresponding hippocampal trace (purple; middle; scale bar: 200 ms) and cross-correlogram of a-IED_HC_ with detected mPFC spindles (lower) during NREM sleep (95% confidence intervals with midpoint represented as black dashed and red lines, respectively; n = 2429 spindles, 3658 stimulations, one sample unkindled rat). **(h)** Spatial representation of clinically identified IED foci (one color per focus) and seizure onset zone (stars; n = 1 sample human subject). Inset demonstrates ratio of IEDs in each focus that are independent (> 1 s apart) from seizure onset zone IEDs to total IEDs in the focus (independent IED ratio). Columns represent seizure onset zone channels and rows represent IED focus channels. **(i)** Histogram of all channels with detected IEDs as classified by independent IED ratio (n = 9 human subjects). Inset demonstrates independent IED ratio across all clinically identified IED foci (n = 9 human subjects).

We hypothesized that emergence of these independent mPFC IEDs was enabled by the repetitive, pathological input received from the hippocampus in the form of hippocampal IEDs. We tested this hypothesis by using repeated (5 s intervals) single pulse stimulation to the hippocampal commissure, a protocol capable of inducing artificial hippocampal IEDs (a-IED_HC_) that resemble spontaneous hippocampal IEDs(*28*) (**Supplementary Figure 1e)**. Application of this protocol to normal rats, in the absence of any kindling, initially resulted in a small, monosynaptic latency evoked response in the mPFC as well as prominent generation of a temporally coupled spindle oscillation (**Figure 1g, Supplementary Figure 1e-f**). Prolonged daily administration of a-IED_HC_ resulted in detection of mPFC IEDs that resembled those identified in kindled rats (**Figure 1g; Supplementary Figure 1g-h**) and occurred in the absence of concurrent spontaneous or artificial hippocampal IEDs. Thus, ongoing occurrence of hippocampal IEDs is sufficient to pathologically modulate the mPFC such that it becomes capable of generating independent IEDs.

These results suggested a link to interictal networks of patients with refractory focal epilepsy, who often display multiple interictal epileptiform foci and IED-spindle coupling(*26*, *29*). We examined sleep intracranial EEG (iEEG) from patients undergoing large-scale electrophysiological monitoring in their presurgical evaluation with multiple IED foci. These subjects expressed 2-9 IED foci, one of which overlapped with the clinically determined seizure onset zone in each subject (**Figure 1h**). IEDs at foci outside of the seizure onset zone had a variable temporal relationship with IEDs in the seizure onset zone, with 0 - 64% co-occurrence (**Figure 1h-i**). Thus, independent IED foci commonly develop in refractory focal human epilepsy.

### Shift in pathological hippocampal-cortical interactions correlates with emergence of independent cortical IED focus

What mediates the emergence of this independent interictal epileptic activity? We established that hippocampal IEDs can reset the phase of the mPFC slow oscillation and induce precisely timed spindles(*10*, *24*, *26*, *27*, *30–32*) (**Figure 1g, Supplementary Figure 2a**). We hypothesized that this pathological oscillatory interaction plays a critical role in the process. Therefore, we examined the relationship between hippocampal IEDs, mPFC delta and mPFC spindle oscillations at a timepoint characterized by sparse independent mPFC IEDs (kindling days 5-10) as compared to a timepoint with frequent, prominent mPFC IEDs (kindling days 15-20). Hippocampal IEDs displayed strong, significant correlation with mPFC spindles (**Figure 2a, upper**), corroborated by a significant increase in spindle power during this epoch (**Supplementary Figure 2b-d**). Unexpectedly, this IED-spindle coordination, though still significant, decreased in magnitude during later kindling stages (**Figure 2a-c**). Across kindling days, there was a progressive diminution of coupled slow oscillations and spindles in the mPFC to incoming hippocampal IEDs (**Figure 2b-c**). Hippocampal IED amplitudes did not decrease during this time (**Supplementary Figure 2e**), suggesting that the plasticity originated from mPFC rather than hippocampus.

**Figure 2.**
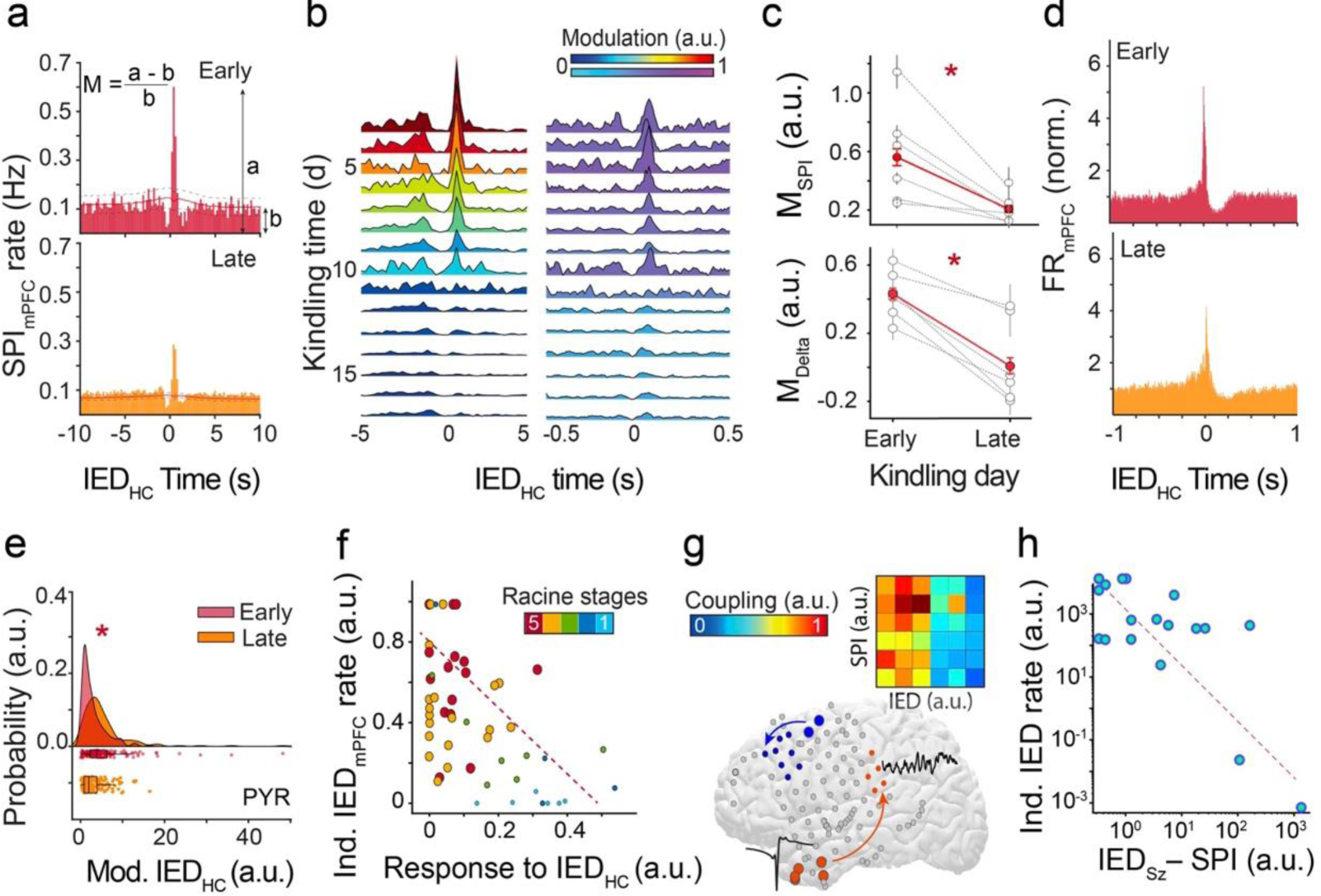
Shift in pathological hippocampal-cortical interactions is related with the emergence of an independent cortical IED focus. **(a)** Sample cross-correlograms of hippocampal IEDs and spindles at early (upper; 1182 IEDs and 5737 spindles from representative rat) and late (lower; 10875 IEDs and 4881 spindles from representative rat) stages of kindling; 95% confidence intervals with midpoint represented as black dashed and red lines, respectively. Parameter *a* represents the peak of the CCG at maximum modulation time and *b* represents the average baseline expected value at this time; coupling modulation (M) is calculated as (a-b)/b. **(b)** Longitudinal modulation of hippocampal IED-spindle (left) and hippocampal IED-slow oscillation (right) coupling across kindling from a representative rat. **(c)** Decrease in coupling modulation from early to late stage of kindling, for hippocampal IED-spindle coupling (upper; n = 6 rats, t = 5.74, P = 9.39 ×10 ^-7^) and hippocampal IED-slow oscillation (SO) coupling (lower; n = 6 rats, t = 7.41, P = 1.31 × 10 ^-9^). **(d)** Averaged peri-event time histogram of clustered mPFC neurons at the time of hippocampal IEDs for early (upper; n = 69 neurons from one sample rat) and late stages of kindling (lower; n = 110 neurons from one sample rat). **(e)** IFR peak probability distribution for clustered mPFC neurons at the time of hippocampal IEDs during early (red) and late stages (orange) of the kindling for putative pyramidal neurons (t = 5.28, P = 1.00 × 10 ^-7^, n = 359 neurons). **(f)** Relationship between responsiveness to hippocampal IEDs (composite scored based on coupling of IED_HC_ with SPI_mPFC_, SO_mPFC_ and MUA_mPFC_) and rate of independent mPFC IEDs (n = 57 sessions from 6 rats; color code indicates Racine stage with dashed lines separating focal from bilateral convulsive seizures; solid black line shows correlation (pairwise linear correlation; P = 2.80 × 10 ^-5^). **(g)** Spatial representation of IED-SPI coupling for two sample IED foci (orange and blue; large circles are IED focus and small circles are maximal region of SPI coupling) from one sample human subject. Inset demonstrates amount of significant IED-SPI coupling occurring between each IED focus; each column represents an IED focus channel (n = 6 IED foci from one sample human subject). **(h)** Relationship between strength of SPI coupling modulation to seizure onset zone IEDs and rate of IEDs independent from those at seizure onset zone (n = 9 human subjects). Coupling strengths and IED rates are normalized within subjects (Spearman r = 0.78; P = 3.30 × 10 ^-7^).

To investigate this notion, we analyzed single neuron activity in the mPFC, classifying clustered units as putative pyramidal cells and interneurons based on established waveform and firing properties (n = 2733 units from n = 6 rats) (*33*). Hippocampal IEDs initially induced a strong increase in mPFC neural spiking at milliseconds latency, but this responsiveness significantly decreased at the later timepoint (**Figure 2d**). This decrease was predominantly mediated by diminished pyramidal cell spiking (**Figure 2e**), though the temporal precision of both hippocampal IED-generated pyramidal cell and interneuron spiking was also reduced (**Supplementary Figure 3, 4a-c**). In parallel, mPFC pyramidal cells and interneurons increased their spiking during independent mPFC IEDs across kindling (**Supplementary Figure 4e-f**). Furthermore, we found that there was a significant negative correlation between mPFC oscillatory and neural spiking responsiveness to hippocampal IEDs and expression of independent mPFC IEDs across rats. These features were additionally able to differentiate the capacity of the epileptic network to generate focal vs. bilateral convulsive seizures (**Figure 2f; Supplementary** Figure 5). Together, these results suggest that mPFC adaptively modulates its activity patterns as hippocampal epilepsy progresses, downregulating output triggered by the epileptic focus but enhancing local interictal epileptic activity.

We next asked whether a similar phenomenon could occur in human subjects with focal epilepsy. Because large-scale, long duration iEEG monitoring is not typically available in human subjects, we examined the relationship between IED-spindle coupling and expression of independent IEDs across the subjects’ multiple clinical IED foci. Clinical IED foci were associated with distinct spatiotemporal patterns of IED-spindle coupling that variably involved other IED foci (**Figure 2g**). Thus, for each clinical IED focus, we quantified the degree to which this brain region expressed spindles temporally coupled to IEDs in the seizure onset zone and the rate of IEDs occurring independently from those in the seizure onset zone. We found a strong negative correlation between these two measures (**Figure 2h**). These results suggest that oscillatory coupling responsiveness decreases as IED foci become more independent from the seizure onset zone.

### Hippocampal IEDs induce hypersynchronous intra-cortical activity patterns

We sought to understand the mechanisms of IED-induced modulation at the neuronal level. In kindled rats, we studied how mPFC neurons respond to a hippocampal IED relative to a comparable physiological epoch. Because hippocampal IEDs reset the cortical slow oscillation phase before coupled spindle induction (**Supplementary Figure 2a**), the resulting pattern resembles the transition from one cortical ‘UP’ state (UP_pre_), through a cortical ‘DOWN’ state, to a subsequent ‘UP’ state (UP_post_; **Figure 3a**). We identified cortical ‘DOWN’ to ‘UP’ state transitions and classified them as either (1) pathological (triggered by a hippocampal IED in UP_pre_ within 200 ms of the ‘DOWN’ state) or (2) physiological (absence of a hippocampal IED) (**Figure 3a-b**). Pathological ‘UP’ states (UP_pre_ and UP_post_) were characterized by increased mPFC population spiking compared to physiological ‘UP’ states, and the pathological ‘DOWN’ state had a more profound decrease in spiking (**Figure 3c**). These firing properties occurred in pyramidal cells and interneurons, (**Supplementary Figure 6**), suggesting that hippocampal IEDs can induce hypersynchronous mPFC spiking that extends hundreds of milliseconds beyond the initial hippocampal-cortical synaptic interaction.

**Figure 3.**
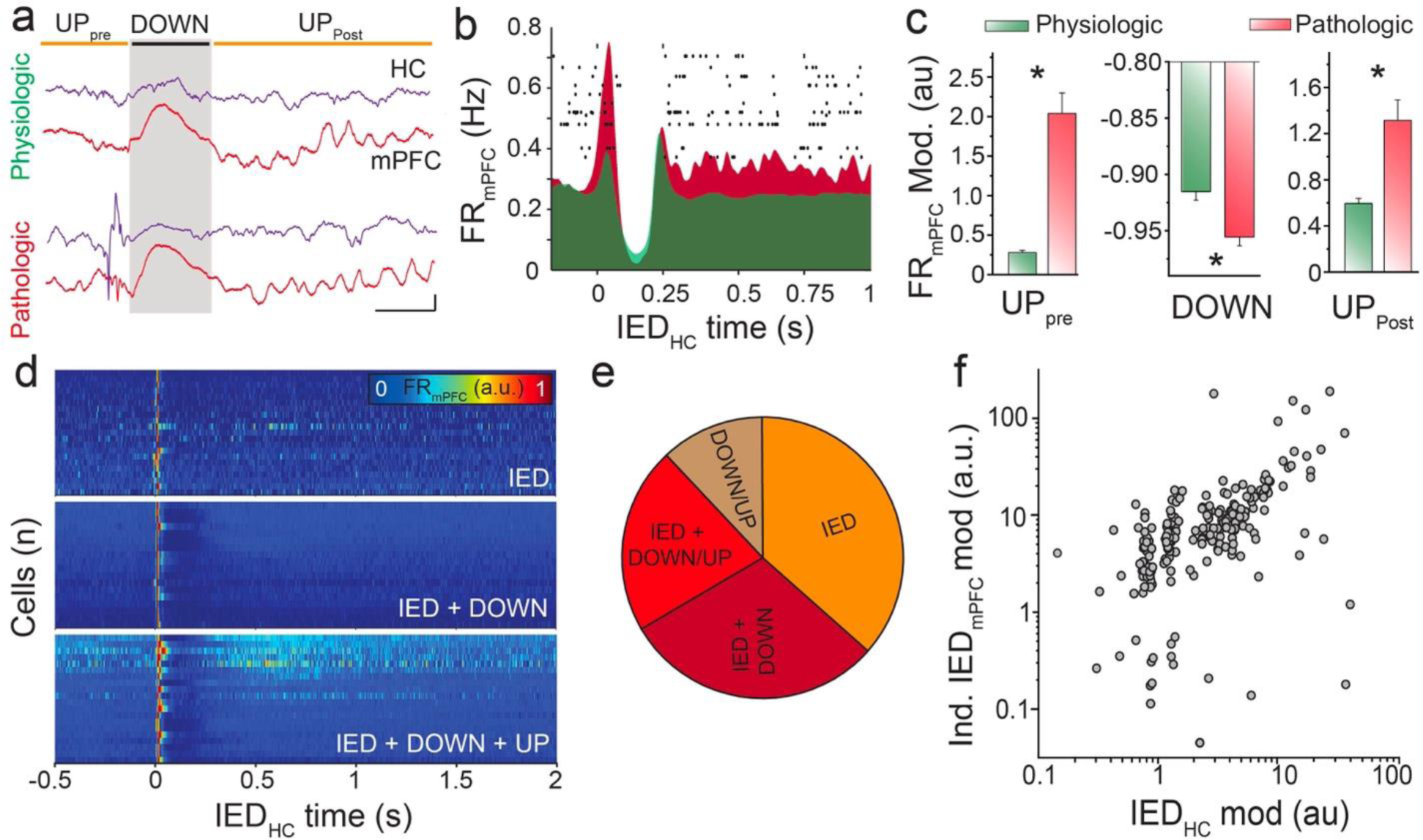
Hypersynchronous mPFC neuronal recruitment into pathological oscillatory sequences primes generation of independent mPFC IED-related neural spiking. **(a)** Sample raw LFP traces from hippocampus (HC, purple) and mPFC (red) demonstrating physiological (upper) and pathological (lower) transition through cortical ‘UP’ and ‘DOWN’ states (scale bar: 500 μV, 200 ms). **(b)** Representative raster plot of mPFC neural spiking (n = 13 sample neurons) overlaid on averaged peri-event firing rate histogram of mPFC neurons during physiological (green) and pathological (red) transition through cortical ‘UP’ and ‘DOWN’ states (n = 2733 neurons from 6 rats). **(c)** Differences in mPFC neural firing rate modulation for physiological (green) and pathological (red) transition through cortical ‘UP’ and ‘DOWN’ states: UP_pre_ (left, averaged normalized values for 100 ms preceding cortical ‘DOWN’ state t = 12.65, P = 9.27 × 10 ^-27^), DOWN (middle, minimum normalized values for 200 ms following onset of cortical ‘DOWN’ state, t = 3.70, P = 2.65 × 10^-4^), UP_post_ (right, averaged normalized values for 500 ms after peak of cortical ‘DOWN’ state, t = –3.93, P = 1.39 × 10 ^-4^; n = 118 sessions). **(d)** Representative normalized firing rate heatmap for mPFC neurons modulated by i) hippocampal IEDs (top), ii) IEDs and subsequent cortical ‘DOWN’ state (middle), iii) IEDs and subsequent cortical ‘DOWN’ and ‘UP’ states (bottom); n = 20 sample neurons per category. **(e)** Proportion of mPFC neurons modulated by categories of pathological events (n = 584 neurons modulated by pathological events out of total 2733 clustered mPFC neurons from 6 rats. IED only: 36.6%; IED + cortical ‘DOWN’ state: 30.0%; IED + cortical ‘DOWN’ + cortical ‘UP’: 21.5%; cortical ‘DOWN’ + cortical ‘UP’ only: 11.9%). **(f)** Relationship between mPFC neural firing rate modulation during hippocampal IEDs and independent mPFC IEDs (n = 323 neurons from 6 rats; r = 0.53; P = 1 × 10 ^-17^).

We next investigated the firing patterns of individual mPFC neurons during these epochs. 21.4% of mPFC neurons exhibited significant firing rate modulation with incoming hippocampal IEDs, and of these, most participated in the evoked pathological DOWN and UP states (**Figure 3d-e**). Only a small proportion of cells was modulated by the cortical DOWN or UP state without any hippocampal IED-related modulation, indicating that a core group of hippocampal-responsive mPFC neurons was responsible for the prolonged hypersynchronous mPFC population patterns. Further, this neuronal core was involved in generating independent mPFC IEDs. There was a significant correlation between strength of hippocampal IED- and independent mPFC IED-driven firing rate modulation across individual neurons (**Figure 3f**). The dual participation of these mPFC neurons suggests a relationship between hippocampal-responsive patterns and the origin of independent mPFC IEDs.

### Spatiotemporally targeted closed-loop mPFC stimulation prevents hypersynchronous cortical response to hippocampal IEDs

We designed a closed-loop electrical stimulation protocol aimed at preventing the hypersynchronous mPFC network response to hippocampal IEDs. We implanted rats with detection electrodes spanning the layers of hippocampal CA1 and stimulating electrodes capable of delivering bipolar electrical stimulation across layers of mPFC. Additional recording electrodes were present in mPFC to monitor response to stimulation. These electrodes were integrated into an embedded system-based responsive device that was set for detection of hippocampal IEDs and delivery of responsive mPFC stimulation (**Figure 4a**). Hippocampal IEDs were identified with sensitivity and specificity similar to conventional offline detection protocols(*34*, *35*) (**Supplementary Figure 7a**). mPFC stimulation waveform was guided by evidence that spindles can be blocked by applying gaussian waves across cortical layers, suggesting desynchronization of the mPFC network(*34*). Closed-loop hippocampal IED-triggered mPFC stimulation resulted in a brief epoch (< 1 s) characterized by a paucity of oscillatory waveforms in the physiological frequency band followed by a recovery of pre-stimulation activity patterns (**Figure 4a-c**). This closed-loop stimulation strongly inhibited spindles, as quantified by significant decrease in spindle band power and IED-spindle cross correlation compared to unstimulated rats (**Figure 4b-d**).

**Figure 4.**
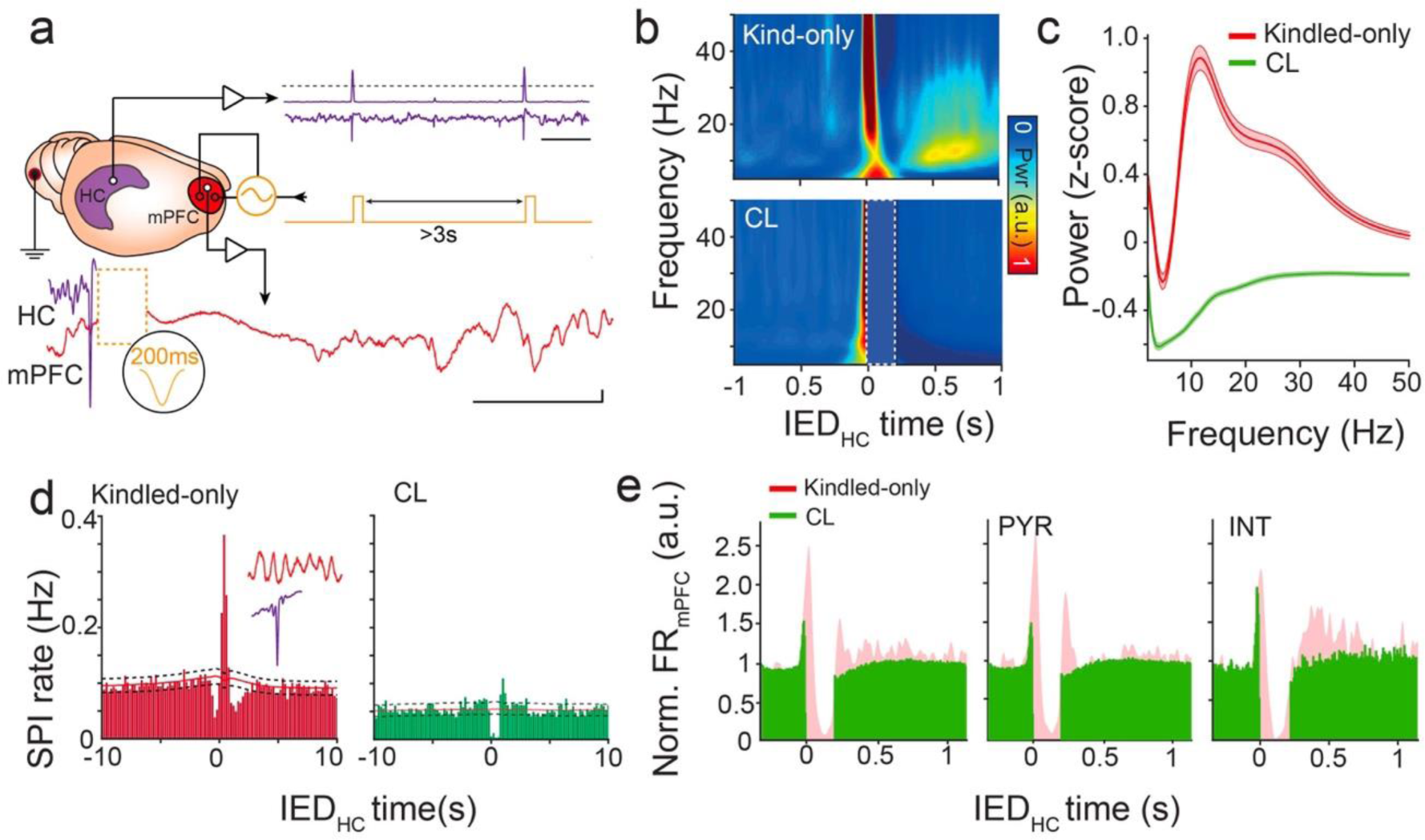
Stimulation of mPFC triggered on hippocampal IEDs prevents pathological oscillatory coupling and neural spiking patterns. **(a)** Schematic of closed-loop intervention demonstrating hippocampal IED detection (purple; scale bar: 1 s), responsively delivered gaussian waveform (yellow), and refractory period between subsequent stimulations (red). Sample raw mPFC LFP trace (red) after closed-loop stimulation triggered on detected hippocampal IED (purple; yellow dashed box represents epoch corresponding to the cropped stimulation artifact; scale bar = 200 μV, 500 ms). **(b)** Sample averaged spectrogram at the time of hippocampal IEDs for a kindled-only rat (upper; n = 1000 randomly sampled IEDs) and a closed-loop stimulated rat (lower; CL; n = 1000 randomly sampled IEDs). Dotted box shows epoch corresponding to the cropped stimulation artifact. **(c)** Sample power spectrum of mPFC LFP following hippocampal IEDs (500 ms interval) in a kindled-only rat (red) and closed-loop stimulated rat (green; n = 1000 IEDs from sample rat for each condition). **(d)** Sample cross-correlogram of hippocampal IEDs with mPFC spindles in a kindled-only rat (left; 8223 IEDs and 9974 spindles) and closed-loop stimulated rat (CL, right; 6687 IEDs and 5475 spindles). 95% confidence intervals with midpoint represented as black dashed and red lines, respectively. **(e)** Averaged peri-event firing rate histogram of mPFC neurons at the time of hippocampal IEDs for closed-loop stimulated rats (green) and kindled-only rats (red), for all neurons (left, CL = 714 neurons from 7 rats; kindled-only = 2733 neurons from 6 rats), putative pyramidal cells (middle, CL = 587 neurons from 7 rats; kindled-only = 2159 neurons from 6 rats) and putative interneurons (right, CL = 127 neurons from 7 rats; kindled only = 574 neurons from 6 rats).

We examined how our closed-loop stimulation protocol affected mPFC neural spiking patterns. Gaussian stimulation resulted in a significant decrease in subsequent population neural spiking compared to the ‘UP’ state following an unstimulated hippocampal IED (**Figure 4e, left**). This change was mediated by both pyramidal cells and interneurons, with pyramidal cells additionally exhibiting a transient decrement in neural spiking relative to baseline (**Figure 4e, middle-right**). Together, these results indicate that hippocampal IED-triggered closed-loop gaussian stimulation can eliminate IED-spindle coupling by decreasing population neural firing across neural subtypes.

### Hippocampal IED-triggered mPFC stimulation prevents expansion of the epileptic network and ameliorates spatial long-term memory deficits

Given the effectiveness of the closed-loop stimulation protocol in decoupling the mPFC from pathological hippocampal input, we investigated its network level and behavioral outcomes. We studied three rat cohorts: i) kindled-only; ii) sham stimulation; and iii) closed-loop stimulation. All animals underwent the same kindling protocol (see **Methods**). The closed-loop stimulation cohort received hippocampal IED-triggered mPFC gaussian stimulation for ∼7 hours after each daily kindling session. The sham stimulation consisted of identical stimulation waveform properties and mean number of stimulations over the same daily duration delivered independently of hippocampal IED timing (**Supplementary Figure 7b**). This stimulation did not consistently change mPFC activity, though mPFC spindle activity was low in the epoch following stimulation (**Supplementary Figure 7c-e**). Neither closed-loop nor sham stimulation caused behavioral state change or altered NREM sleep characteristics, but small changes in REM sleep activity were observed, consistent with involvement of mPFC in REM regulation(*36*); **Supplementary Figure 8**).

Rats that underwent the closed-loop stimulation protocol exhibited significantly decreased development of mPFC IEDs compared to kindled-only and sham stimulated animals (**Figure 5a-b; Supplementary Figure 9a**). This reduction was progressive over the course of kindling, with the most profound differences observable during the late phase (**Figure 5a-b**). To further investigate interictal epileptogenicity of the mPFC, we designed an index that quantifies the sharpness and predictability of the LFP by examining its temporal first derivative (**Supplementary Figure 9b**). The closed-loop stimulation protocol was selectively effective in preventing an increase in interictal epileptogenicity (**Figure 5c; Supplementary Figure 9c**). In parallel, we observed that rats undergoing the closed-loop stimulation protocol did not develop bilateral convulsive seizures during a duration of kindling that robustly generated this seizure semiology in control kindled rats. Open-loop stimulation delayed, but did not prevent, this progression (**Figure 5d, Supplementary Figure 10a**). These results suggest that closed-loop gaussian mPFC stimulation triggered on hippocampal IEDs inhibits cortical recruitment into the mesial temporal epileptic network and preserves physiological properties of mPFC LFP.

**Figure 5.**
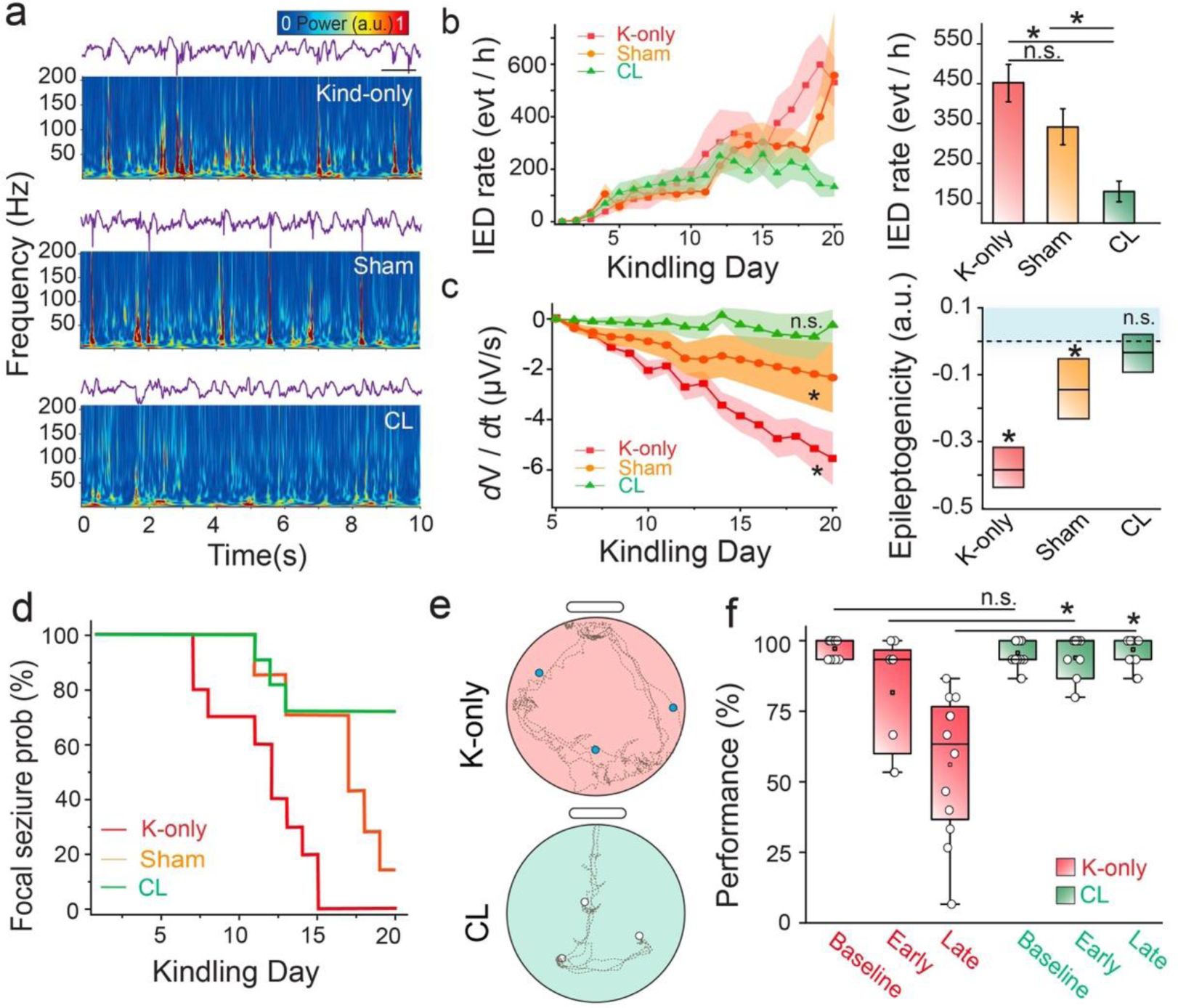
Closed-loop stimulation prevents epilepsy progression and memory deterioration. **(a)** Raw LFP traces from hippocampus (purple) and simultaneously acquired mPFC spectrogram for sample kindled-only rat (upper), sham stimulated rat (middle) and closed-loop stimulated rat (lower; scale bar: 1s). **(b)** Occurrence of mPFC IEDs across kindling (left) with quantification of mPFC IED rate during late kindling days (days 15 – 20; right); ANOVA with Bonferroni-Holm correction, P = 2.93 × 10^-6^, F = 25.47; kindled-only vs. sham, P = 0.3287; closed-loop vs. kindled-only; P = 2.64 × 10^-6^; closed-loop vs. sham, P = 0.0049; kindled-only n = 10 rats; sham n = 7 rats; closed-loop n = 11 rats). **(c)** Epileptogenicity analysis of mPFC LFP in NREM sleep over kindling progression (left; Mann-Kendall tau; closed-loop, n = 11 rats, P = 0.3; sham, n = 7 rats, P = 8.43 × 10^-8^; kindled-only, n = 10 rats, P = 8.42 × 10^-8^). Regression of the epilepsy progression over kindling days (right; linear mixed effects model; closed-loop, P = 0.18836; sham, P = 0.001; kindled-only, P = 3.5128 × 10^-26^). **(d)** Survival curve for progression from focal to bilateral convulsive seizures based on Racine stages (kindled-only; n = 10 rats; sham stimulated, n = 7 rats; closed-loop stimulated, n = 11 rats). **(e)** Long-term spatial memory consolidation test. Representative examples of exploration path (dashed lines) with reward locations (blue circle, non-retrieved reward; open circle, retrieved reward) for the first three trials of the memory test in the late kindling stage for sample kindled-only and closed-loop stimulated rat. **(f)** Memory test performance (percentage of rewards retrieved during the first three trials) over kindling for kindled-only (n = 4 rats) and closed-loop stimulated rats (n = 6 rats): ANOVA with Bonferroni-Holm correction: F = 20.27; baseline (P = 0.7487, n = 12 kindled-only and 17 closed-loop stimulated sessions), early kindling (P = 0.0407, n = 8 kindled-only and 11 closed-loop stimulated sessions) and late kindling (P = 1.72 **×** 10 ^-12^, n = 12 kindled-only and 17 closed-loop stimulated sessions).

We used a cheeseboard maze(*37*, *38*) to assay the effect of our closed-loop stimulation protocol on this cognitive process. The behavioral protocol, consisting of a training session followed by a testing session the following day, was conducted prior to initiation of kindling, every 5 days during the kindling, and at the conclusion of the kindling. Rats were trained on the spatial location of three hidden water rewards, and all animals were able to demonstrate effective learning by the end of the training session. On behavior training days, closed-loop stimulation was initiated after the completion of training and continued for approximately 7 hours post-training (**Supplementary Figure 10b**). Kindled-only animals displayed an early and progressive decrement of long-term memory performance as assayed by ability to retrieve water rewards during the test trials. In contrast, rats who additionally underwent closed-loop stimulation maintained baseline high levels of memory performance (**Figure 5e-f**). Thus, this closed-loop network intervention is also capable of preserving long-term memory capacity, indicating an overall protective effect on both mPFC network activity and function.

## Discussion

We demonstrate that temporally specific inhibition of pathological hippocampal-cortical oscillatory coupling prevents recruitment of synaptically connected cortex into the epileptic network and preserves long-term memory in a rodent focal epilepsy model. This closed-loop electrical stimulation intervention decreased hypersynchronous cortical neural spiking, blocking the prolonged and amplified response associated with uninterrupted IED-spindle coupling. IED-spindle coupling occurs in children and adults with focal epilepsy(*24*, *25*, *34*, *39*), and we found a similar relationship between oscillatory coupling and independent IED foci in patients with epilepsy.

We found that interictal activity can initiate a process that over time enlarges the brain territory capable of independently generating IEDs. When patients express IEDs outside of the seizure onset zone, the degree of IED independence varies, potentially indicating a similar longitudinal process. Epilepsy surgery has an increased likelihood of favorable outcome when the entire irritative zone is addressed, suggesting that impeding expansion of this zone could have therapeutic implications(*40*). However, our inability to define the network organization of the irritative zone limits its utility in planning surgical resection and determining causal relationships to cognitive comorbidities(*41–43*). Our results support that measures which inhibit development of independent IED foci can prevent long-term memory deficits.

In our rodent model, the mPFC responded strongly and consistently to hippocampal IEDs by mounting an exaggerated version of the physiological reaction to hippocampal output(*44*): a burst of cortical neural spiking followed by a ‘DOWN’ state and spindle oscillation (during the following ‘UP’ state). This responsiveness decreased as kindling progressed, paralleled by an increase in capacity for local hypersynchronous neural spiking and epileptic activity. In patients, a similar tradeoff revealed decreased IED-spindling coupling modulation associated with increased incidence of independent IEDs. Our cellular level observations suggest a pathological adaptation in epilepsy(*45*), and could underpin the interictal network segregation identified by resting state functional MRI in patients with progressive focal epilepsy(*46*).

The closed-loop electrical stimulation protocol we employed was designed and tested to eliminate pathological hippocampal-cortical coupling by preventing expression of a cortical ‘DOWN’ state and subsequent sleep spindle when delivered in response to a hippocampal IED. We affirmed the efficacy of this stimulation in eliminating IED-spindle coupling and further determined that it prevented the synchronized increase in population firing associated with the uninterrupted response of the mPFC network to hippocampal IEDs. When provided on an ongoing basis, such stimulation was capable of preventing establishment of independent mPFC IEDs and normalizing mPFC network parameters, despite concurrent daily hippocampal seizures. These results suggest that hippocampal input in isolation is insufficient to drive long-lasting cortical network change; the subsequent cortical response is an integral contributor, potentially by instantiating local plasticity processes. Open loop stimulation, which induced a similar transient decrease in population neural firing, was capable of delaying seizure progression, but was ultimately less effective in modulating mPFC network activity than the closed-loop stimulation, in keeping with the notion that temporally targeted approaches can more sustainably modify networks(*47*).

Because closed-loop electrical stimulation is clinically used in patients with epilepsy(*48*, *49*), our approach has high translational potential: (1) it does not require viral vectors or genetic modifications; (2) the safety of electrical stimulation is established and (3) technologic advancements enhance computational capacity and decrease invasiveness of electronic devices(*35*, *50*). Here, we were limited in our ability to characterize the temporal determinants of the stimulation, including the duration of efficacy after stimulation was stopped, and whether a similar efficacy could be obtained if stimulation was initiated within an established epileptic network. Our findings could apply to clinical situations to prevent epileptogenesis after brain insult,(*51*, *52*) and support further investigation.

Finely tuned and highly regulated hippocampal-cortical communication is required for multiple cognitive processes. Our results emphasize that hippocampal-cortical dynamics during the interictal state are modifiable targets to ameliorate epilepsy-associated memory dysfunction. We provide evidence for plasticity that is instantiated by chronic interictal epileptic activity patterns and eventually downregulates hippocampal input, at the expense of increased local mPFC hypersynchrony. Rebalancing the hippocampal-cortical interaction by inhibiting the pathological mPFC response may prevent this plasticity and preserve physiological mPFC activity patterns needed for memory consolidation. Thus, spatiotemporally targeted interventions that block the network effect of IEDs may modify disease course and ameliorate cognitive comorbidities in individuals with focal epilepsy.

## Methods

### Animal surgery procedure

All animal experiments were approved by the Institutional Animal Care and Use Committee at Columbia University Irving Medical Center. Thirty male and female Long Evans rats (200 to 350 g) were used for intracranial implantation. Rats were kept on a regular 12h-12h light-dark cycle and housed in pairs prior to implantation but separated afterward. Prior experimentation was not performed on these animals. The animals were initially anesthetized with 2% isoflurane and maintained under anesthesia with 0.75 to 1% isoflurane during surgery. Silicon probes (Neuronexus) and/or 50 μm diameter tungsten wires mounted on custom-made micro-drives were implanted in the hippocampus (−3.5 anterior-posterior (AP) and 3.0 medial-lateral (ML)) and ipsilateral mPFC (3.5 AP, 0.2 ML, -2.5 DV). Closed-loop and sham stimulated rats were implanted with bipolar stimulation electrodes (50 μm diameter tungsten wires separated by 500 μm) spanning across mPFC cortical layers and in line with the recording electrodes. A pair of stimulating electrodes (two 50 μm diameter tungsten wires attached together with 500 μm dorsoventral tip separation) was implanted into the hippocampal commissure (-0.5 AP, 0.8 ML, - 4.2 DV) for electrical kindling stimulation. Screws in the skull, overlying the cerebellum, served as ground electrodes. The craniotomies were covered by Gelfoam and sealed using a 10:1 mixture of paraffin and mineral oil. Rats recovered for 4 to 5 days prior to initiation of experimentation. Hippocampal electrodes were adjusted in the dorsal-ventral axis to span the layers of CA1 based on localizing neurophysiological signals.

### Kindling stimulation

Kindling stimulation consisted of 2 s duration bipolar current pulses (60 Hz, 1 ms pulse width) and was delivered twice per day (20 min separation interval between stimulations). The amount of current used was determined on the initial kindling days by titrating current in 5 μA increments starting at 25 μA (10 min separation interval between stimulations) until a hippocampal seizure greater than 20 s duration was generated. This current setting was used for the remainder of kindling. 5-7 hours of post-kindling electrophysiological recordings were performed.

### Closed-loop stimulation

Rats underwent the kindling procedure as previously described. Closed-loop stimulation consisted of 5 - 8 V gaussian waves of 200 ms duration that were delivered across cortical layers. Stimulation was adjusted to create gaussian waves opposite in polarity to cortical delta waves and voltage was titrated to the minimum required to suppress spindles. Two rats had misplacement of mPFC stimulation electrodes and were removed from further experimentation. Starting 1h after the second seizure induction, closed-loop stimulation commenced and was maintained for approximately 7 hours per day.

### Sham stimulation

Stimulation was performed as described above, but cortical electrical stimulation was not coupled to the online detection of hippocampal IEDs. The frequency of stimulations corresponded to the average number of stimulations of closed-loop treated rats at the equivalent day of the kindling procedure.

### Artificial hippocampal IEDs

Square pulses (200 μs) delivered to the hippocampal commissure were used for induction of artificial IEDs. The stimulation voltage was adjusted to elicit hippocampal IEDs of amplitude comparable to that of spontaneous hippocampal IEDs in kindled rats (0.5 to 2.5 mV). Artificial IEDs were elicited every 3-5 s for 6 - 12 hr per day.

### Neurophysiological data acquisition and closed-loop system

Neurophysiological signals were amplified, digitized continuously at 20 kHz using a head-stage directly attached to the probe (RHD2000 Intan technology), and stored for off-line analysis with 16-bit format. For real-time IED detection and closed-loop stimulation, hippocampal data were routed via RHD2000 digital to analog converter into a custom 32-bit microcontroller (STM32)-based module. The data were filtered using an active bandpass filter (60 to 80 Hz). Subsequently, the filtered data were rectified and convolved with a moving average window to generate the instantaneous power of the signal at the frequency band of interest. The instantaneous power was then compared to the noise threshold, which was defined based on the standard deviation of 10 s of baseline, filtered data. Surpassing the threshold triggered delivery of the preprogrammed cortical stimulation using a stimulus generator (STG4002; Multichannel Systems). Stimulation times were digitized and stored for off-line analysis. To avoid subsequent delivery of stimulation during the period of network response, a refractory period for stimulation of 3 s was put in place. System parameters were visualized online via a graphical user interface, allowing for on-demand noise floor and stimulation threshold adjustment. Additionally, a 3-axis accelerometer signal from the animal’s head-stage amplifier was continuously analyzed using a MATLAB custom algorithm to prevent stimulation triggered by mechanical, movement-related artifacts.

### LFP preprocessing

Data were analyzed using MATLAB (MathWorks) and visualized using Neuroscope. The electrophysiological data were resampled to 1250 Hz to facilitate LFP analysis. Epochs of sleep were identified by immobility in the motion signal of the animal’s onboard accelerometer and absence of electromyogram (EMG) artifacts. NREM and REM sleep epochs were classified using a validated automated sleep scoring algorithm based primarily on ratios of cortical delta (0.5 – 4 Hz) and hippocampal theta (5 – 8 Hz) (*53*). Sleep-scoring was visually inspected and manually adjusted if necessary using whitened spectrograms and raw traces. Activity patterns were detected using custom MATLAB code based on the Freely Moving Animal (http://fmatoolbox.sourceforge.net) toolbox during NREM sleep epochs.

### Racine stages

Seizures induced by kindling were monitored by an overhead video camera. The severity of the seizures was scored according to Racine stages: 1, mouth and facial movements; 2, head nodding; 3, forelimb clonus; 4, rearing with forelimb clonus; 5, rearing and falling with forelimb clonus. Stages 4 and 5 were considered as development of bilateral convulsive seizures.

### Human subjects

We analyzed intracranial electroencephalography (iEEG) recordings from 9 patients with focal epilepsy who underwent clinical electrode placement as part of the work-up for epilepsy surgery. The Institutional Review Board at New York University Langone Medical Center (NYULMC) approved gathering and analysis of this data. Informed written consent was obtained from all patients. Patients were eligible if they were diagnosed with focal epilepsy, had continuous high quality iEEG recordings, lacked major cortical lesions, and had > 1 clinically identified IED focus.

### Clinical reports

Clinical iEEG reports were obtained for each patient’s hospital admission, detailing localization of IEDs and the clinically identified seizure onset zone. Clinical interpretation was performed using a combination of referential montage (referenced to epidural electrodes) and bipolar montage (based on pairs of neighboring electrodes).

### iEEG data pre-processing and detections

Epochs of sleep were analyzed, and these were identified by immobility on synchronized video in concert with increased delta/gamma frequency ratio in the iEEG spectrogram. Referential data was imported into MATLAB and resampled from 512 to 1250 Hz for compatibility with previously validated analytical toolboxes. IED and spindle detection were performed on all electrodes as previously defined, in addition to IED-IED and IED-SPI coupling metrics(*24*). Channels with IED-SPI coupling modulation less than 0.001 were excluded from analysis.

### iEEG electrode localization

MNI coordinates of electrodes were determined by reconstruction of subject specific pial surfaces, co-registration of pre- and post-implant MRI images, a combination of manual and automatic localization of electrodes, and subsequent co-registration to a standard template brain(*54*).

### IED detection

IEDs were detected by: (i) band-pass filtering at 50-85 Hz and signal rectification; (ii) detection of events for which the filtered envelope surpassed the median filtered signal by at least 5 standard deviations; (iii) elimination of events for which the waveform amplitude (high pass filtered above 15 Hz) did not surpass the mean baseline signal by at least 10 standard deviations; (iv) elimination of events for which the waveform amplitude surpassed the mean baseline by over 100 standard deviations (consistent with artifact). Independence of IEDs was defined as the absence of a co-occurring IED in another brain region within 100 ms.

### Spindle detection

Cortical LFP was filtered between 10-20 Hz using a Butterworth filter. The filtered signal was then rectified, and instantaneous power was extracted using the Hilbert transform. Spindles were detected when the filtered envelope was at least 2 standard deviations above the filtered baseline with an interposed peak at least 4 standard deviations, but not more than 14 standard deviations, above this baseline. The filtered baseline standard deviation was calculated after epochs of cortical IEDs were removed. Additionally, spindle duration was defined as 350 – 3000 ms, with detected events occurring within 250 ms merged into a single event.

### Cortical DOWN state detection

DOWN states were detected based on identification of large positive deflections in the cortical LFP that were associated with decreases in the multiunit activity firing rate(*55*). First, cortical LFP was filtered (0.5–6 Hz) and subsequently z-scored which yielded *Z* (t). Next, the start (t_start_), peak (t_peak_), and end (t_end_) of putative DOWN states were defined as upward-downward-upward zero-crossings of the derivative of Z(t). Events with Z(t_peak_) > 1 and Z(t_end_) < −1.5 or Z(t_peak_) > 2 and Z(t_end_) < 0, were deemed candidate events. Lastly, events with > 500 ms or <150 ms durations were discarded. LFP-based DOWN state detection was validated by the instantaneous mPFC multiunit activity. Events where the multiunit activity decreased relative to t_peak_ were considered DOWN states. All detections were visually inspected for accuracy for each recording session. Pathological DOWN/UP transitions were classified as those that initiated within 200 ms of hippocampal IEDs. The remaining transitions were classified as physiological. The phase of the DOWN state in the delta band was derived using Hilbert transformation of the filtered signal. For closed-loop and sham stimulated rats, the stimulation artifacts (200 ms) were removed from the recordings before performing the corresponding event detections.

### Time domain cross-correlograms and coupling modulation

To determine coupling between detected oscillations, cross-correlograms (CCGs) were calculated using a modified convolution method, as previously described(*20*, *26*, *56*). 95% confidence intervals were estimated from a Poisson distribution with the mean lambda value determined from the convolution. The peak of the CCG (a) above the 95% confidence interval and expected baseline value level (b) at time zero enabled calculation of the coupling modulation (M) as a normalized ratio: M = (a-b)/b.

### Frequency domain analysis

Spectrograms were generated using an analytical wavelet transformation (Gabor). Spindle band power was extracted from z-scored NREM power spectra.

### LFP epileptogenicity

To examine changes to NREM waveforms across kindling in a detection-free manner, segments of NREM sleep (duration = 5 s; n = 10 segments per session) were selected using a random number generator and with manual verification. Each segment was passed through a Savitzky-Golay filter to equalize the content of higher frequencies across segments. The gradient of each filtered sample was taken and values greater than a noise-level determined by a median-based threshold were removed. These threshold values were normalized by z-scoring across kindling days and were fit to a linear regression model to derive the slope LFP changes and their statistical significance.

### Spiking-data processing

Noise-free epochs of data were used to first generate realistic neural spike templates and perform noise floor estimation. Multiunit activity was detected on the basis of spike amplitude using derivative-and-shift peak finding and median-based thresholding methods. Spike sorting was performed on bandpass filtered (250 – 2500 Hz) data using KiloSort(*57*). Manual cluster cutting and curation to segregate single neurons were performed using Phy (https://github.com/kwikteam/ Phy). Single-units were further validated based on observation of mean waveform shape, auto-correlogram, and consistency of the localization of the mean waveform with the probe geometry. Putative excitatory and inhibitory neurons were identified based on their auto-correlograms and waveform characteristics using CellExplorer(*33*).

### Unit modulation and instantaneous firing rate

Zenith of Event-based Time-locked Anomalies (ZETA) was used to determine whether neurons show a statistically significant time-dependent firing rate modulation relative to an event in a manner that avoids arbitrary parameter selection and binning(*58*). ZETA identified neurons that showed significantly modulated spiking activity (p < 0.05) with respect to hippocampal and mPFC detected events, and yielded instantaneous firing rate (IFR) amplitudes and latencies. Raw IFR peak values were normalized by the baseline firing rate of each cell to quantify the normalized IFR modulation of each cell with respect to a reference event. IFR latency was computed by computing the temporal lag between the reference event and IFR peak. Recording sessions containing less than 20 events were excluded from analysis. Firing modulation (M) was calculated as (a-b)/b, defining *a* as average firing for the time interval of interest, and *b* as the baseline firing, which was calculated by averaging the firing rate over 500 ms of baseline activity. To ensure that observed single unit activity measures were not driven by variability across animals, a generalized linear mixed-effects model was employed with rat identity as a random effects term(*59*). Population rates were computed by summing all the detected single unit activity with 1 ms resolution and smoothing the resulting population rate vector with a 50 ms gaussian window.

### Cheeseboard maze memory test

Rats were placed on a water deprivation schedule for 3 to 5 d prior to intracranial implantation to ensure they could receive water through a handheld syringe. Rats were weighed daily during water deprivation to ensure that body weight did not decrease to <85% of pre-deprivation measurements. Behavior for all tasks was tested on a cheeseboard maze as previously described(*24*).

Prior to surgery, water-deprived rats were first familiarized with exploring the maze environment to obtain water. Initially, the rat was placed in the center of the maze and allowed to explore and retrieve multiple (∼25) randomly placed hidden water rewards. Over the next 3 d, the number of available water rewards on the maze was gradually reduced, and a trial structure was introduced such that the rat received a food reward (0.5 to 1 Froot Loop) after successful retrieval of all water rewards. The rat was then trained to return to the starting box after retrieving water to obtain its food reward. After 2 to 4 d of this repeated procedure, the rat would consistently explore the maze to obtain three spatially distinct water rewards and then independently return to the starting box. To prevent use of odor-mediated searching, the maze was wiped with a towel soaked in 70% ethanol and rotated by a random multiple of 90° relative to the starting box between all trials.

Each memory cycle on this task was completed over 2 days. On the first day, the rats learned the location of three hidden water rewards placed in a randomly selected set of 3 water wells over the course of ∼40 trials (25 trial sessions, then ∼3-h home cage rest, and then 15 trial sessions). All rats obtained >90% performance averaged over five trials by the end of the training session. On the second day, the rat was given a three-trial test with water rewards located in the same location as the first day to assess memory for the spatial configuration of the reward locations.

Memory performance in the test session was scored by determining the percentage of rewards obtained per trial. Behavior sessions were monitored by an overhead video camera and tracking of the rat’s location was facilitated by blue and red light emitting diodes (LEDs) attached to its cap. Rats were trained to obtain > 80% memory performance in the test session prior to intracranial electrode implantation. After the implantation and recovery, the rats were tested in three memory cycles before starting the kindling procedure to establish baseline memory performance. Subsequently, the rats were tested every 5 days during the kindling. Learning trials started at least 5 hr after seizure induction and test trials were performed prior to the subsequent day’s kindling. Closed-loop electrical stimulation was performed in the home cage after behavioral sessions.

### Histology

After completion of experimentation, placement of the implanted electrodes was verified for all the rats. Rats were euthanized with sodium pentobarbital and perfused via the heart with saline phosphate buffer (PBS) followed by 4% paraformaldehyde (PFA). Whole brains were extracted and post-fixated 48 hr in 4% PFA, embedded in agar 5% and sectioned using a VT-1000S vibratome (Leica) to obtain 70 μm coronal slices. Slices were permeabilized in PBS with Triton X-100 0.25%, stained with DAPI (1:10000) for 20 min, and mounted with Fluoromount-G (Invitrogen). A fluorescence Revolve microscope (ECHO) was used to image the slices. Tungsten wires or probe location were reconstructed from adjacent slices.

### Statistics

Statistical analysis was performed using a combination of freely available, online MATLAB toolboxes (Freely Moving Animal Toolbox; http://fmatoolbox.sourceforge.net), custom MATLAB code and Origin Pro. Differences between groups were calculated using paired t-test, or ANOVA (Bonferroni-Holm correction) depending on the nature of the data analyzed. Mann-Kendall tau was used to identify significant longitudinal trends. Error bars represent mean ± standard error of the mean (SEM). Significance level was P < 0.05.

## Acknowledgments

This work was supported by Columbia University Irving Medical Center, Department of Neurology, and Columbia University, School of Engineering and Applied Science. This work was supported by the National Institute of Health grants R01NS118091, R21 EY 32381-01, RF1NS128669, National Science Foundation 1944415 and 2219891. We would like to thank all Khodagholy and Gelinas laboratory members for their support.

## Author contributions

DK and JNG conceived the project. JJF, LM, TK, and JE performed animal experimentation. AH, JJF, ZY, CW, SS, DK, and JNG performed data analysis. ZZ, and JJF fabricated devices and optimized the closed-loop stimulation system. WD was the attending neurosurgeon. O.D. supervised the human epilepsy recordings and processes related to the institutional review board. All authors contributed to writing the paper.

## Data and materials availability

All data needed to evaluate the conclusions in the paper are present in the paper and/or the Supplementary Materials. Additional data related to this paper may be requested from the authors.

