## Supplementary material for "Closed-loop electrical stimulation to prevent focal epilepsy progression and long-term memory impairment": SI

†These authors contributed equally.

**The PDF file includes:**

Figs. S1 to S10

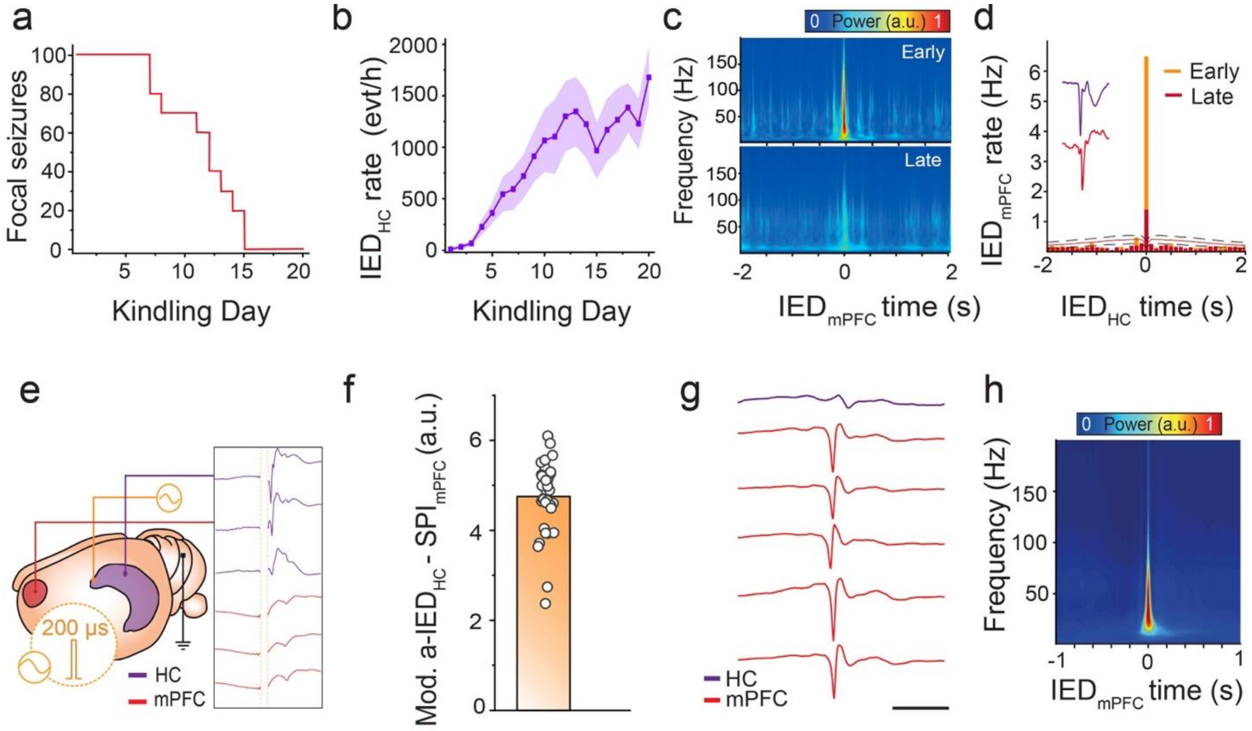


**Supplementary Figure 1: Kindling protocol and artificial IED induction.**

**(a)** Survival curve for progression from focal to bilateral convulsive seizures based on Racine stages (n = 10 rats).

**(b)** Occurrence of hippocampal IEDs over kindling (n = 10 rats; shaded error bars represent SEM).

**(c)** Averaged normalized spectrogram of hippocampal activity at the time of mPFC IEDs for early (upper) and late (lower) stages of kindling.

**(d)** Cross-correlograms of hippocampal and mPFC IEDs at early (orange; 557 IEDs and 11980 spindles) and late stages (red; 1469 IEDs and 9026 spindles) of the kindling from sample rat; 95% confidence intervals with midpoint represented as black dashed and red lines, respectively.

**(e)** Schematic of hippocampal commissure pulse stimulation (200 µs). Representative hippocampal LFP trace demonstrating artificial IED (a-$IED_{HC}$) and mPFC LFP trace showing evoked response.

**(f)** Coupling modulation values (M = (a-b)/b) derived from cross-correlograms of a-$IED_{HC}$ and mPFC spindles (n = 35 sessions, from 2 rats).

**(g)** Induction of a-$IED_{HC}$ leads to development of independent mPFC IEDs in the absence of kindling. Averaged LFP traces (left) showing independent mPFC IED waveform across mPFC recording sites (red) in absence of hippocampal IED (purple; scale bar = 500 ms).

**(h)** Induction of a-$IED_{HC}$ leads to development of independent mPFC IEDs in the absence of kindling. Average spectrogram of independent mPFC IEDs (n = 680 IEDs from one sample unkindled rat).

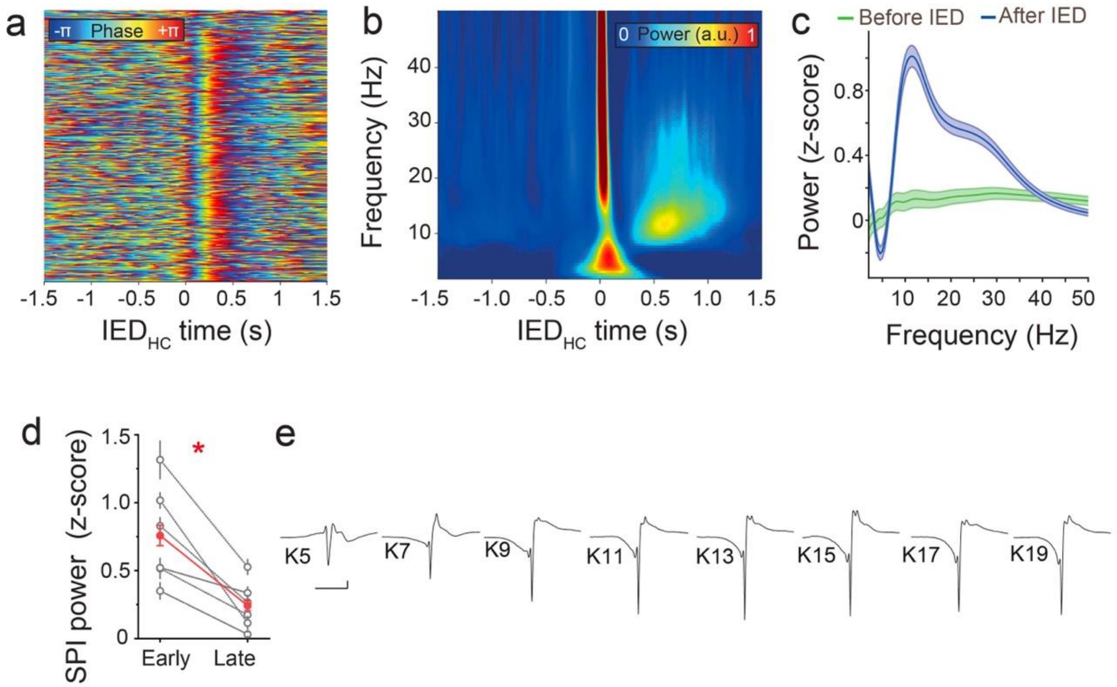


**Supplementary Figure 2: Modulation of IED-spindle coupling across kindling.**

**(a)** Stacked trials of mPFC delta phase (2-5 Hz) aligned to detection of hippocampal IED (blue = -pi, red = pi, n = 414 trials in one sample rat).

**(b)** Sample trigger-averaged spectrogram of mPFC power in early kindling (n = 1000 IEDs).

**(c)** Sample power spectrum before (green; -1000 to -500 ms interval) and after (blue; 500 to 1000 ms interval) for early kindling (n = 1000 randomly selected IEDs in one rat; shaded bars are SEM).

**(d)** Decrease in spindle power from early to late stage of kindling across rats (n = 6 rats, t = 6.17, P = 5.44 $\times 10^{-7}$).

**(e)** Average recorded hippocampal IED waveforms over kindling days (K; n = 500 randomly selected IEDs for each session in one rat; scale bar = 200 µV, 100 µs).

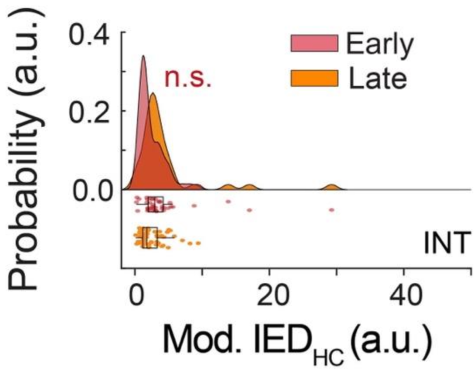


**Supplementary Figure 3: mPFC interneuron spiking responses to hippocampal IEDs.**

IFR peak probability distribution for clustered mPFC neurons at the time of hippocampal IEDs during early (red) and late stages (orange) of the kindling for interneurons ($t = 1.93$, $P = 0.057$, $n = 101$ neurons).

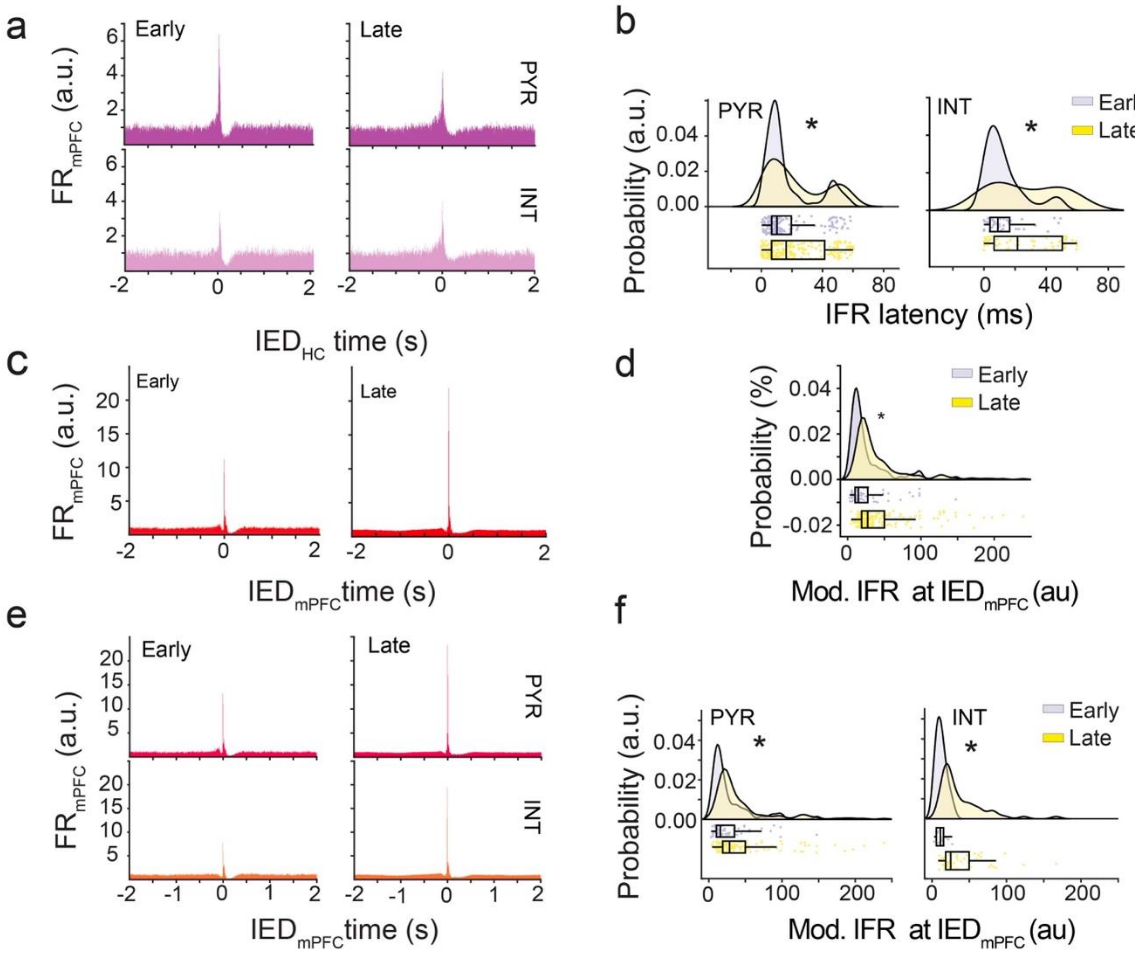


**Supplementary Figure 4: mPFC neural spiking responses to hippocampal and mPFC IEDs.**

**(a)** Averaged peri-event firing rate histogram of mPFC neurons at the time of hippocampal IEDs for early (left) and late (right) stage of kindling, for pyramidal neurons (upper) and interneurons (lower).

**(b)** mPFC neuron IFR latency probability distribution at the time of HC IEDs in early (grey) and late stages (yellow) of the kindling, for pyramidal neurons (upper, $t = -2.91$, $P = 0.0039$, $n = 359$ neurons), and interneurons (lower, $t = -3.68$, $P = 0.00037$, $n = 101$ neurons).

**(c)** Averaged peri-event firing rate histogram of mPFC neurons at the time of independent mPFC IEDs for early (left) and late stage of kindling (right).

**(d)** mPFC neural population IFR probability distribution at the time of independent mPFC IEDs in early (grey) and late stages (yellow) of the kindling $t = -2.11$, $P = 0.030$, $n = 268$ neurons).

**(e)** Averaged peri-event firing rate histogram of mPFC neurons at the time of independent mPFC IEDs in early (left) and late (right) stage of kindling, for pyramidal neurons (upper) and interneurons (lower).

**(f)** mPFC neuron IFR probability distribution at the time of independent mPFC IEDs in early (grey) and late (yellow) stages of kindling, for pyramidal neurons (upper, $t = -2.30$, $P = 0.020$, $n = 195$ neurons), and interneurons (lower, $t = -3.21$, $P = 0.0020$, $n = 73$ neurons).

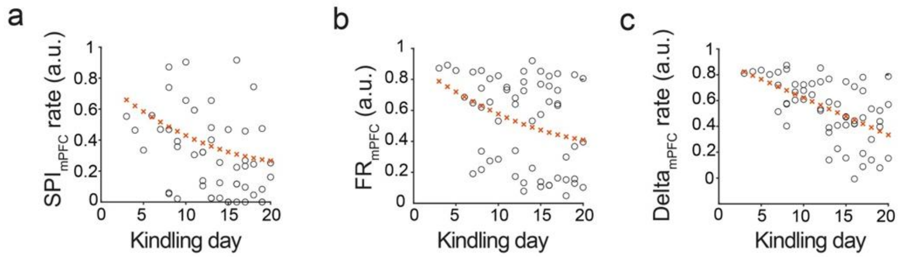


**Supplementary Figure 5: mPFC oscillatory and neural spiking responsiveness decreases over kindling.**

**(a)** Hippocampal IED-mPFC spindle coupling modulation decreases across kindling (n = 57 sessions from 6 rats).

**(b)** Hippocampal IED-evoked mPFC neural firing decreases across kindling (n = 57 sessions from 6 rats).

**(c)** Hippocampal IED-mPFC slow oscillation coupling modulation decreases across kindling (n = 57 sessions from 6 rats).

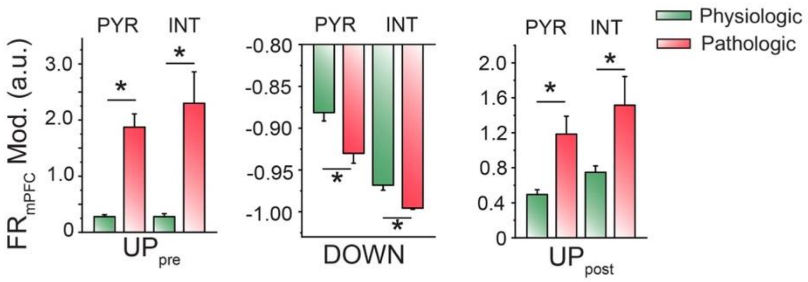


**Supplementary Figure 6: mPFC pyramidal cell and interneuron firing through cortical 'UP' and 'DOWN' states.**

Differences in mPFC pyramidal cell and interneuron firing rate modulation for physiological (green) and pathological (red) transition through cortical 'UP' and 'DOWN' states: $UP_{pre}$ (left, averaged normalized values for 100 ms preceding cortical 'DOWN' state, pyramidal cells: $P = 1.22 \times 10^{-16}$; $t = 10.16$; interneurons: $3.14 \times 10^{-12}$, $t = 8.14$), DOWN (middle, minimum normalized values for 200 ms following onset of cortical 'DOWN' state, pyramidal cells: $P = 0.0021$; $t = 3.14$; interneurons: $P = 4.52 \times 10^{-5}$, $t = 4.47$), $UP_{post}$ (right, averaged normalized values for 500 ms after peak of cortical 'DOWN' state, pyramidal cells: $P = 0.0016$, $t = -3.27$; interneurons: $P = 0.026$, $t = -2.29$; $n = 118$ sessions).

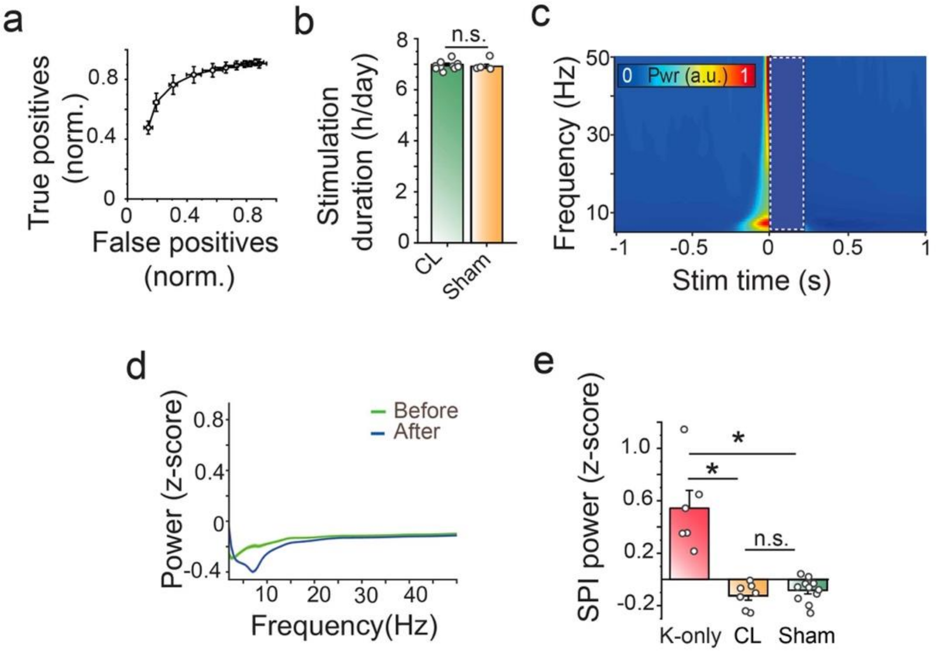


**Supplementary Figure 7: Properties of closed-loop stimulation protocol.**

**(a)** Receiver operating characteristics curve for the online IED detection compared to the offline detection (n = 9946 IEDs, 5 sessions from 5 rats).

**(b)** Duration of stimulation treatment per day is equivalent between closed-loop (CL, n = 10 rats; green) and sham stimulated rats (n = 7 rats; blue); t = 0.69; P = 0.5017.

**(c)** Averaged mPFC spectrogram at the time of sham stimulation (n = 1000 randomly selected stimulations in one sample rat).

**(d)** Averaged mPFC power spectrum before (green, calculated from -1000 to -500 ms interval) and after (blue, 500 to 1000 ms interval) for sham stimulation (n = 1000 randomly selected stimulations in one sample rat).

**(e)** Change in mPFC spindle band power from before to after hippocampal IED in kindled-only rats (n = 6 rats), sham stimulation (n = 7 rats; $P = 9.69 \times 10^{-7}$ vs. kindled-only) and closed-loop stimulation (n = 11 rats; $P = 1.51 \times 10^{-6}$ vs. kindled-only; P = 0.6539 vs. sham stimulation). ANOVA with Bonferroni-Holm correction, F = 28.38.

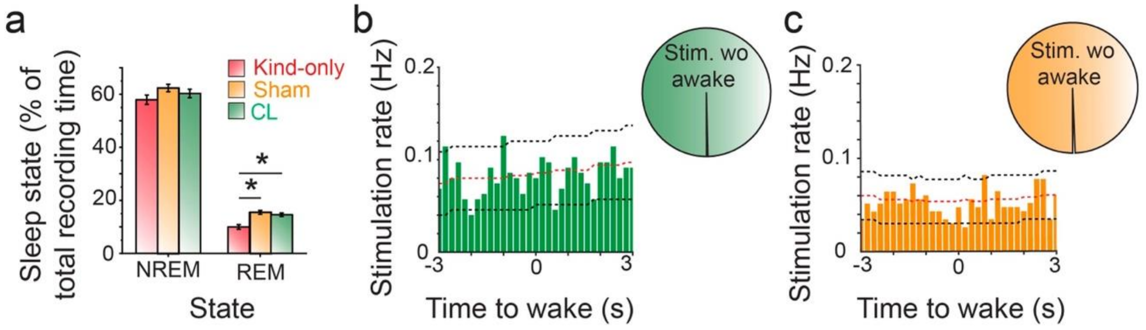


**Supplementary Figure 8: mPFC electrical stimulation does not alter NREM sleep architecture or induce state change.**

**(a)** NREM and REM sleep time as a percentage of the total duration of recording sessions. NREM: ANOVA with Bonferroni-Holm correction F = 2.54; kindling-only vs. sham, P = 0.2333; kindling-only vs. closed-loop, P = 0.2347; sham vs. closed-loop, P = 0.5562; REM: ANOVA with Bonferroni-Holm correction F = 11.11; kindling-only vs. sham, P = $3.05 \times 10^{-4}$; kindling-only vs. closed-loop, P = 0.0038; sham vs. closed-loop, P = 0.5453.

**(b)** Cross-correlogram of mPFC electrical stimulations with transitions to wake state for closed-loop stimulation; 95% confidence intervals with midpoint represented as black dashed and red lines, respectively. Pie chart shows the percentage of stimulations not related to transitions to wake state (99.7% of 22280 stimulations, 876 wake transitions).

**(c)** Cross-correlogram of mPFC electrical stimulations with transitions to wake state for sham stimulation; 95% confidence intervals with midpoint represented as black dashed and red lines, respectively. Pie chart shows the percentage of stimulations not related to transitions to wake state (99.4% of 20397 stimulations, 1166 wake transitions).

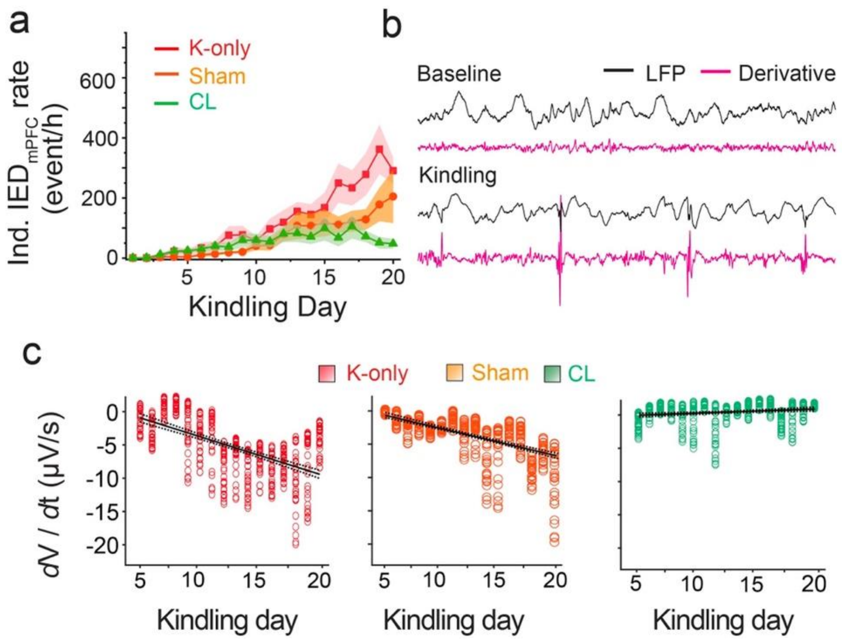


**Supplementary Figure 9: Decrease in mPFC epileptogenicity with closed-loop stimulation.**

**(a)** Occurrence of independent mPFC IEDs in mPFC in NREM sleep over kindling progression.

**(b)** Sample first derivative trace (magenta) of wide-band mPFC LFP trace (black) prior to kindling (upper) and in kindled (lower) states.

**(c)** Variation in first derivative across kindling for sample kindled-only rat, sham stimulated rat, and closed-loop stimulated rat.

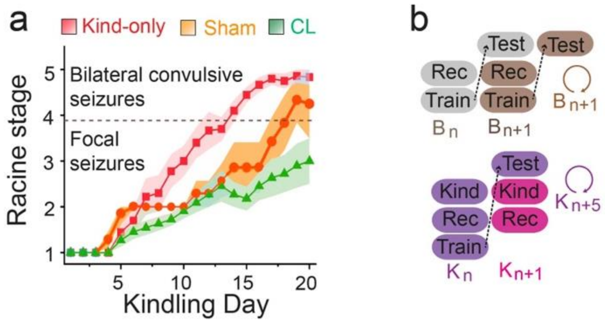


**Supplementary Figure 10: Seizure progression and behavioral protocol.**

**(a)** Racine stage progression over kindling for kindled-only rats (n = 10 rats), sham stimulated rats (n = 7 rats) and closed-loop stimulated rats (CL, n = 11 rats).

**(b)** Schematic of behavioral protocol during baseline (B, upper) and kindling (K, lower), showing the successive sessions during the memory training and testing: delivery of kindling stimulation (Kind), neurophysiological recording in absence or presence of closed-loop stimulation (Rec), behavioral training (Train), and behavioral testing the subsequent day (Test).
